# E3 ligase ASB3 downregulates antiviral innate immunity by targeting MAVS for ubiquitin-proteasomal degradation

**DOI:** 10.1101/2023.12.10.570965

**Authors:** Mingyang Cheng, Yiyuan Lu, Jiarui Wang, Haixu Wang, Yu Sun, Wenhui Zhao, Junhong Wang, Chunwei Shi, Jiawei Luo, Ming Gao, Tianxin Yu, Jianzhong Wang, Jiayao Guan, Nan Wang, Wentao Yang, Yanlong Jiang, Haibin Huang, Guilian Yang, Xin Cao, Dongqin Yang, Chunfeng Wang, Yan Zeng

## Abstract

E3 ubiquitin ligases are very important to regulate antiviral immunity during viral infection. Here, we discovered that Ankyrin repeat and SOCS box-containing protein 3 (ASB3), an E3 ligase, are upregulated in the presence of RNA viruses, particularly Influenza A virus (IAV). Notably, overexpression of ASB3 inhibits type I IFN (IFN-I) responses induced by Sendai virus (SeV) and H9N2, and ablation of ASB3 restores SeV and H9N2 infection-mediated transcription of IFN-β and its downstream interferon-stimulated genes (ISGs). Interestingly, animals lacking ASB3 showed a decreased susceptibility to H9N2 and PR8 infections. Mechanistically, ASB3 interacts with MAVS and directly mediates K48-linked polyubiquitination and degradation of MAVS at K297, thereby inhibiting the phosphorylation levels of TBK1 and IRF3, downregulating downstream antiviral signaling. These findings establish ASB3 as a critical negative regulator in controlling the activation of antiviral signaling and describe a novel function of ASB3 that has not been previously reported.

**IMPORTANCE:** IAV is a significant zoonotic pathogen that causes infections of the respiratory system. Hosts have evolved multiple strategies to defend against IAV infection. However, not all host proteins play an active defense role. In this study, we found that the E3 ligase ASB3 regulates antiviral immunity by manipulating MAVS stability. Briefly, overexpression of ASB3 degrades MAVS, thereby promoting viral replication. In contrast, ASB3 deletion restores MAVS expression, upregulating IFN-I responses. Additional research revealed that ASB3 mediates the K48-linked polyubiquitination of MAVS at K297, resulting in ASB3 being degraded via the ubiquitin-proteasome pathway. These findings reveal, for the first time, a novel mechanism by which ASB3 negatively regulates antiviral immunity and provides a potential target for anti-IAV drug development.

## INTRODUCTION

The innate immune system serves as the host’s first line of defense against viral infections, with Type I interferons (IFN-I) considered a critical component of the antiviral innate immunity (1). Viral nucleic acids are recognized by pattern recognition receptors (PRRs), which activate downstream signaling cascades. RIG-I-like receptors (RLRs), such as retinoic acid-inducible gene I (RIG-I)-like receptors and melanoma differentiation-associated gene 5 (MDA5), are primarily responsible for recognizing viral RNA (2). In contrast, viral DNA is sensed by cyclic GMP-AMP synthase (cGAS) (3). Mitochondrial antiviral signaling proteins (MAVS) or stimulator of interferon genes (STING) is activated as adapter molecules and initiate signaling, which drives the recruitment of downstream TANK-binding kinase 1 (TBK1)/I-kappa B kinase ε (IKKε) (4). This cascade then causes nuclear factor-κB (NF-κB) and interferon regulatory factor 3 (IRF3) to be phosphorylated, facilitating their translocation into the nucleus and causing the synthesis of IFN-I (5, 6). Influenza A virus (IAV) is a major zoonotic pathogen that poses a considerable challenge to influenza management and containment due to its broad host tropism and frequent genetic segment mutation and reassortment (7). IAV is a negative-stranded RNA virus belonging to the *Orthomyxoviridae* family. Numerous studies have demonstrated that IAV infection is capable of limiting the host’s innate immune responses (8), and infection in mice was exacerbated by the H5 subtype of highly pathogenic avian influenza virus with the I283M/K526R mutation on PB2 (9). Our previous study also showed that PB1-mediated degradation of MAVS promoted H7N9 virus infection (10). In fact, throughout infection, the virus and host interact dynamically, and the host has the power to control the IFN-I response in a variety of ways.

A growing body of research suggests that hosts use several cellular proteins to shield MAVS and trigger effective antiviral immune responses. For instance, TRIM31 facilitates MAVS aggregation and activation through lys63-linked polyubiquitination (11), and UBL7 enhances antiviral innate immunity by promoting lys27-linked polyubiquitination of MAVS (12). However, not all proteins trigger defense mechanisms to protect the host during viral infection. According to recent studies, NOG1 targets IRF3 binding to the promoter, inhibiting type I IFN production (13), and SIRT2 disrupts the formation of the G3BP1-cGAS complex by deacetylating G3BP1, inhibiting cGAS activity and activation of downstream signaling pathways (14). Therefore, understanding how host proteins regulate immune responses is essential for effective viral clearance.

Protein ubiquitination modifications intricately govern the regulatory dynamics within the RLR signaling pathway. Ubiquitination as a 76 amino acid small molecule protein requires three levels of enzymatic reactions to link the lysine on the substrate, including the E1 enzyme for activation, the E2 enzyme for delivery, and the E3 enzyme for ligation (15). Ankyrin repeat and SOCS box-containing protein 3 (ASB3) are members of the ASB family. Unlike other E3 enzymes, the ASB protein requires binding to three auxiliary proteins (Rbx2, Cullin 5, and Elongin B/C) in order to form an E3 enzyme complex that catalyzes ubiquitination (16). A previous study has documented that the E3 enzyme ASB1, belonging to the same ASB family, effectively suppresses the TAB2-linked K48 polyubiquitination, consequently facilitating the initiation of the NF-κB and MAPK signaling cascades (17). Conversely, ASB8 actively promotes the degradation of TBK1/IKKε K48-linked polyubiquitination and exerts a downregulatory influence on IFN-I production (18). The discussion surrounding ASB3 focuses on how it contributes to the ubiquitin-proteasome-dependent destruction of the tumor necrosis factor receptor (TNF-R2) (19). The E3 enzymes known to modulate the signaling of RLRs primarily belong to the TRIM and RNF families. However, no investigations have been discovered concerning the modulation of the RLR signaling pathway by ASB3.

Our study showed that, during IAV infection, the E3 ligase ASB3 functions as a negative regulator of RLR-mediated innate antiviral immunity. Overexpression of ASB3 led to reduced expression levels of IFN-I, while ASB3 deficiency activated TBK1 and IRF3 phosphorylation to trigger IFN-I production, thereby inhibiting the replication of IAV. Mechanistically, ASB3 specifically interacts with MAVS and further blocks the IFN-I response by catalyzing K48-linked polyubiquitination of MAVS at Lys297 and undergoing proteasomal pathway degradation. According to our study, the ASB3 protein plays a new function in the host immune system’s defense against the influenza virus.

## RESULTS

### ASB3 expression may depend on IAV infection

To confirm the key role of ASB3 in viral infection, A549 cells were infected with Sendai virus (SeV), herpes simplex virus 1 (HSV-1), or stimulated with poly(I:C) (an analog of viral dsRNA) and 2’3’-cGAMP (a ligand of STING) for 0, 6, 12, or 18 h, the results showed that the protein levels of ASB3 were significantly elevated in A549 cells infected with SeV or stimulated with poly(I:C) but not HSV-1 and 2’3’-cGAMP compared with the untreated cells (Fig. 1A and B). Given the phenomenon of ASB3 upregulation in RNA virus infection, we next addressed the role of ASB3 in response to IAV, a negative-sense RNA virus with a segmented genome. Here, we choose two representative strains for infection: A/chicken/Hong Kong/G9/1997 (H9N2) and A/Puerto Rico/8/34 (H1N1, PR8). Similarly, ASB3 remained highly expressed in A549 cells infected with H9N2 and PR8 (Fig. 1C and D). Meanwhile, we also observed ASB3 localization in infected cells by confocal microscopy. Fluorescence signals of IAV NP and ASB3 protein were significantly increased after H9N2 or PR8 infection (Fig. 1E and F). Our findings indicate that ASB3 could be induced by IAV infection.

**Fig. 1.**
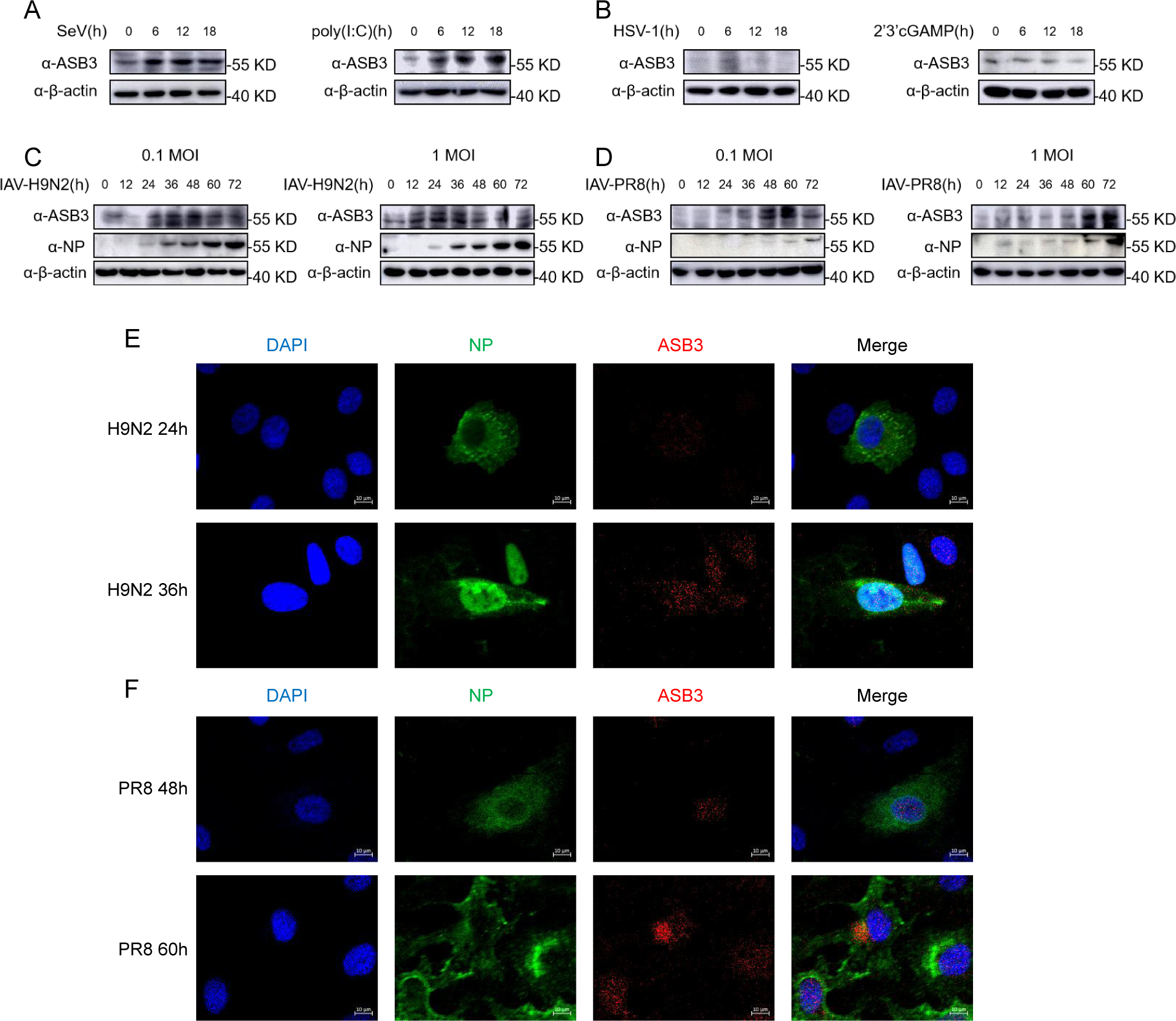
ASB3 expression induced by IAV infection. (A and B) A549 cells were infected with SeV and HSV-1 (MOI=1) or stimulated with poly(I:C) (10 μg/ml) and 2’3’cGAMP (10 μg/ml) for 0, 6, 12, or 18 h. Then, the mRNA and protein levels of ASB3 in A549 cells were detected by qPCR and immunoblot analysis. (C and D) A549 cells were infected with H9N2 or PR8 (0.1 MOI and 1 MOI) for 0, 12, 24, 36, 48, 60, and 72 h, and the protein levels of ASB3 were analyzed by immunoblot analysis. (E and F) A549 cells were infected with H9N2 (1 MOI) or PR8 (1 MOI) for the indicated time. The subcellular localization of endogenous ASB3 and NP was analyzed by fluorescence microscopy.

### ASB3 negatively regulates viral RNA-triggered signaling

We noted that the heightened expression of ASB3 was concomitant with a reduction in the phosphorylation levels of TBK1, IRF3, and IκBα during IAV infection, thereby prompting our speculation that ASB3 might exert a pivotal function in the host’s innate defense against viral infections. To confirm this, HEK-293T cells were transfected with IFN-β-Luc and ISRE-Luc along with HA-ASB3 expression plasmids and infected with SeV or stimulated with poly(I:C). The results showed that overexpression of ASB3 inhibited SeV- and poly(I:C)- (but not cGAS and STING-) induced IFN-β and ISRE promoter activity in a dose-dependent manner (Fig. 2A). To further investigate the influence of the ASB3 protein on the host’s antiviral response, we assessed the gene transcription levels of IFN-β and its downstream IFN-stimulated genes (ISGs) or the phosphorylation levels of signaling molecules in SeV or H9N2 infected HEK-293T cells. We found that overexpression of ASB3 significantly inhibited gene transcription levels of IFNB1, ISG15, IFIT2, and OSAL (Fig. 2B). In addition, we found that the recombinant vesicular stomatitis virus expressing green fluorescent protein (VSV-GFP) infection was significantly promoted in HEK-293T cells transfected with an increasing amount of exogenous ASB3 plasmids (Fig. 2C). We also observed that overexpression of ASB3 exacerbated the compromised phosphorylation of TBK1, IRF3, and IκBα induced by H9N2, which indicated further impairment of the antiviral response (Fig. 2D). Indeed, the attenuation of the innate immune response mediated by ASB3 coincided with an increase in viral replication (Fig. 2E and F). These results indicated that ASB3 is an important negative regulator of viral RNA-mediated antiviral innate immune response.

**Fig. 2.**
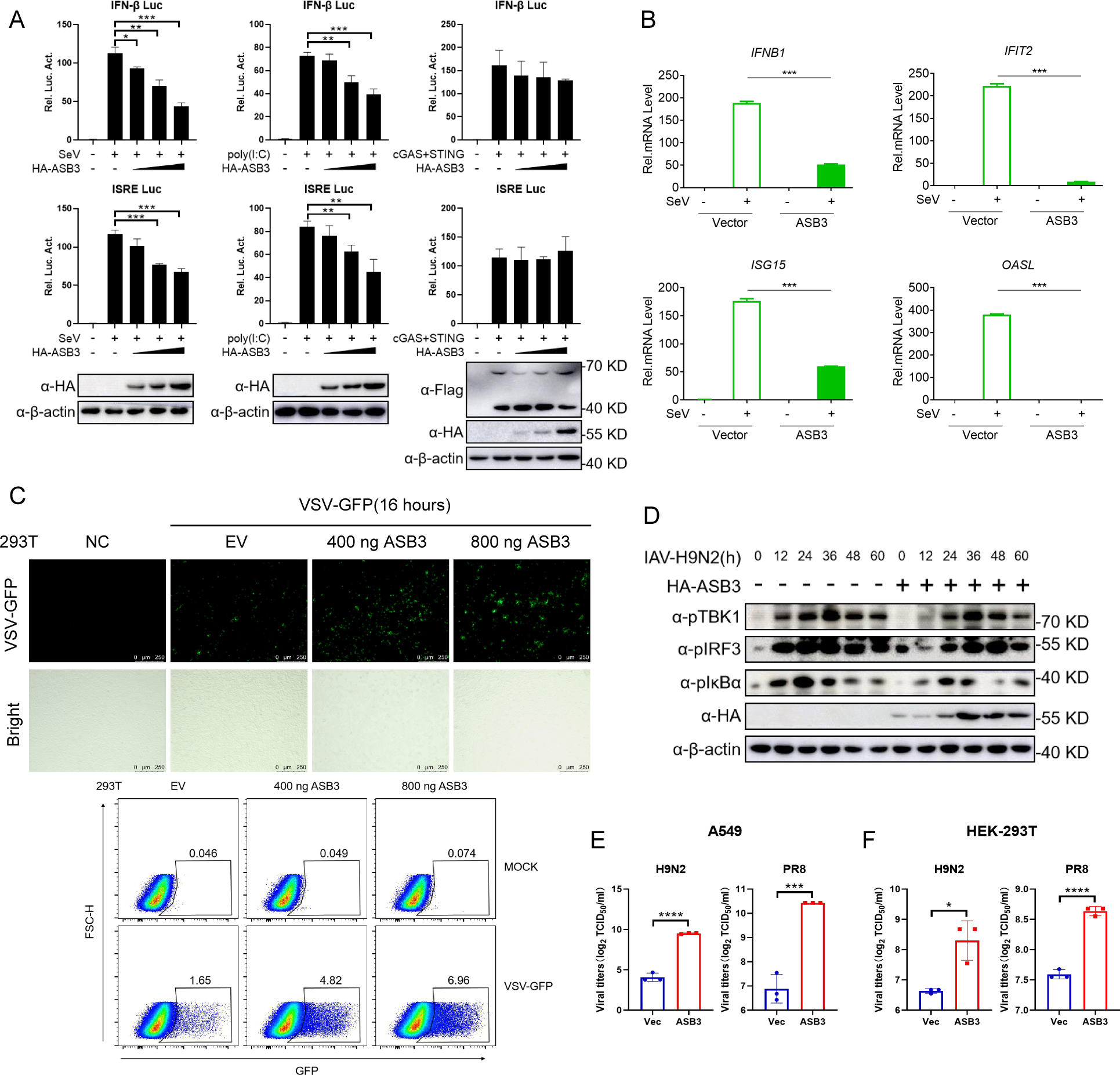
ASB3 negatively regulates type I IFN production. (A) HEK-293T cells were transfected with IFN-β-Luc, ISRE-Luc (100 ng), or Flag-tagged cGAS and STING and pRL-TK plasmid (50 ng) along with increasing HA-ASB3-expressing plasmids (50, 100, 200 ng). After 24 h, indicated cells were infected with SeV (MOI=1) or transfected with poly(I:C) (10 μg/ml). The protein level and promoter activation were determined by the dual-luciferase reporter gene and Western blotting assay. (B) HEK-293T cells were transfected with 1 μg of HA empty vector (EV) or HA-ASB3 expression plasmid. After 24 h, the cells were infected with SeV (MOI=1) for an additional 12 h. The mRNA expression of IFNB1, IFIT2, ISG15, and OASL was measured by qPCR. (C) Fluorescence microscopy analysis and flow cytometric analysis of the replication of VSV-GFP in HEK-293T cells transfected with EV or increasing HA-ASB3 expression plasmid at indicated dose for 16 h, followed by treatment with or without VSV-GFP (MOI=0.1) infection at indicated time. (D) HEK-293T cells were transfected with 1 μg of HA empty vector or HA-ASB3 expression plasmid. After 24 h, the cells were infected with H9N2 (MOI=1) for 0, 12, 24, 36, 48, and 60 h. The protein expression of pTBK1, TBK1, pIRF3, IRF3, pIκBα, IκBα, HA, and β-actin were measured through immunoblot analysis. (E) Viral titers in A549 cells transfected with ASB3 plasmid and infected with H9N2 virus or PR8 virus (n=3). (F) Viral titers in HEK-293T cells transfected with ASB3 plasmid and infected with H9N2 virus or PR8 virus (n=3).

### ASB3 deficiency enhances the cellular antiviral responses

To determine whether endogenous ASB3 is equally capable of negatively regulating the IFN-I response, bone marrow-derived macrophages (BMDM) and primary kidney epithelial cells isolated from ASB3^+/+^ mice and ASB3^-/-^ mice were infected with SeV or H9N2 for 12 h. As expected, ASB3 deficiency significantly promoted SeV-induced gene transcription levels of Ifnb1, Isg56, Cxcl10, Ccl5, IL-6, and TNFα in BMDM (Fig. 3A). Compared with primary kidney epithelial cells from ASB3^+/+^ mice, primary kidney epithelial cells from ASB3^-/-^ mice infected with SeV showed higher gene transcription levels of Ifnb1, Isg15, Isg56, Cxcl10, Mx1, and Oasl1 (Fig. 3B).

**Fig. 3.**
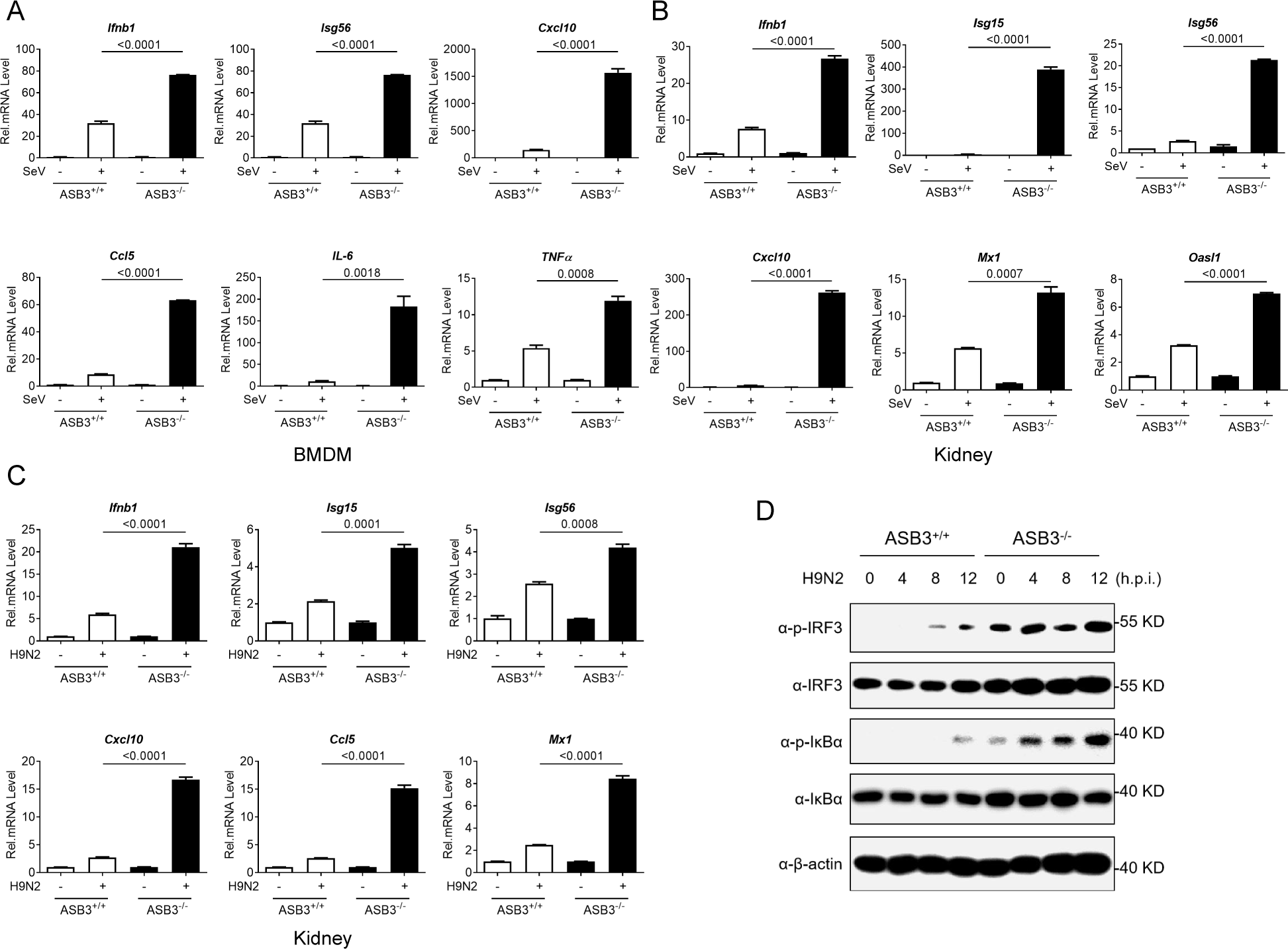
ASB3 deficiency enhances type I IFN response. (A and B) Bone marrow-derived macrophages (BMDM) and primary kidney epithelial cells isolated from ASB3^+/+^ and ASB3^-/-^ mice were infected with SeV (MOI=1) for 12 h. The mRNA expression of Ifnb1, Isg15, Isg56, Cxcl10, Ccl5, Mx1, Oasl, IL-6, and TNFα were measured by qPCR. (C) Primary kidney epithelial cells were infected with H9N2 (MOI=1) for 12 h. The mRNA expression of Ifnb1, Isg15, Isg56, Cxcl10, Ccl5, and Mx1 were measured by qPCR. (D) BMDM were infected with H9N2 (MOI=1) for 0, 4, 8, and 12 h. The protein expression of pIRF3, IRF3, pIκBα, IκBα, and β-actin were measured immunoblot analysis.

Similarly, We also found higher gene transcript levels of Ifnb1, Isg15, Isg56, Cxcl10, Ccl5, and Mx1 in ASB3-deficient primary kidney epithelial cells infected with H9N2 (Fig. 3C). Importantly, ASB3 deficiency enhanced phosphorylation of TBK1, IRF3, and IκBα induced by H9N2 (Fig. 3D). Collectively, these results suggest that ASB3 deletion enhances IAV-induced IFN-I responses.

### ASB3 specifically interacts with MAVS

To identify the regulatory role of ASB3 in the IAV-triggered RLRs-IFN-β signaling axis, we transfected the RIG-IN, MDA5-N, MAVS, or IRF3-5D plasmids together with the IFN-β or ISRE promoter in the presence or absence of ASB3. We observed that overexpression of ASB3 inhibited RIG-IN-, MDA5-N-, and MAVS-induced IFN-β or ISRE promoter activity in a dose-dependent manner, but not TBK1- and IRF3-5D-induced IFN-β or ISRE promoter activity (Fig. 4A and B). Since the inhibition occurs upstream of TBK1, we speculated that ASB3 is strongly associated with MAVS. Further coimmunoprecipitation (Co-IP) experiments revealed that ASB3 interacted specifically with MAVS but not with TRAF3 (Fig. 4C). cGAS and STING serve as critical receptors and junction proteins in the immune response against DNA viruses. Interestingly, our findings indicate that ASB3 does not interact with either cGAS or STING (Fig. 4D). Confocal microscopy confirmed that ASB3 and MAVS (but not TRAF3) co-localized in the cytoplasm (Fig. 4E). *In vitro* transcription-translation assays also showed that ASB3 directly interacted with MAVS (Fig. 4F). Additionally, semi-endogenous and endogenous IP experiments demonstrated an enhanced interaction between ASB3 and MAVS during infection with H9N2 or PR8 virus (Fig. 4G and H). Domain mapping experiments indicated that ASB3 interacted with MAVS through its ANK domain (Figs 4I). In summary, these findings indicate that MAVS represents a specific target of ASB3, further highlighting the distinct role of ASB3 in anti-RNA viral mechanisms.

**Fig. 4.**
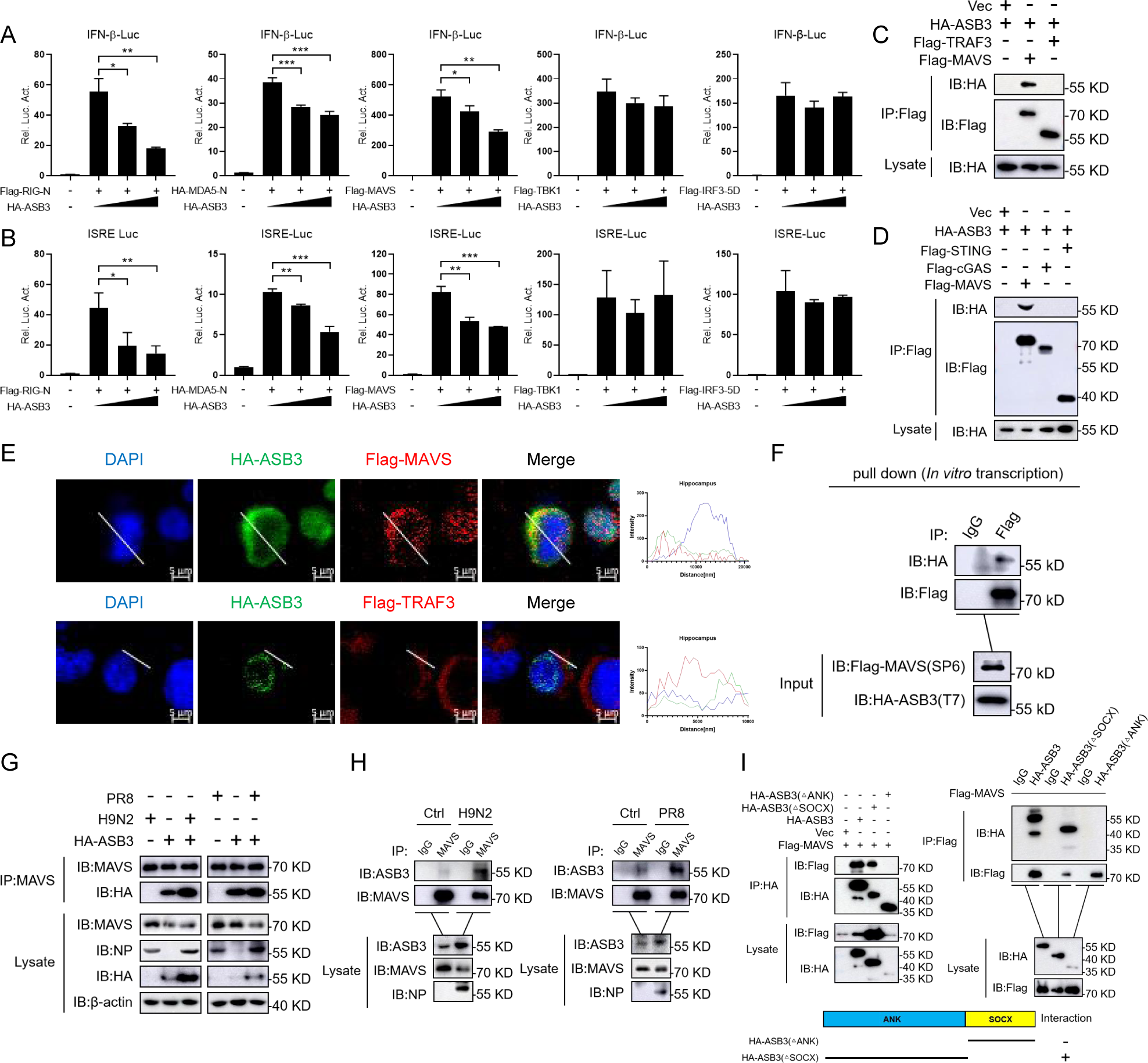
ASB3 specifically interacts with MAVS. (A and B) HEK-293T cells were transfected with the indicated plasmids (RIG-IN, MDA5-N, MAVS, TBK1, and IRF3-5D) along with the control vector or increased amounts of ASB3 expression plasmids. IFN-β and ISRE promoter activity were performed 24 h after transfection. (C and D) Co-IP analysis of the interaction of ASB3 with MAVS in HEK-293T cells transfected indicated plasmids (MAVS, TRAF3, cGAS, STING, and ASB3). (E) HEK-293T cells were transfected with the indicated plasmids (MAVS and TRAF3) along with the ASB3 expression plasmid. Then, it is stained with a Flag antibody or HA antibody and a secondary antibody. DAPI (blue), Flag-MAVS or TRAF3 (red), and HA-ASB3 (green). Scale bars, 5 μm. (F) In vitro transcription pull-down analysis of the interaction of ASB3 with MAVS. (G and H) Semi-endogenous and endogenous Co-IP analysis of the interaction of ASB3 with MAVS in HEK-293T or A549 cells infected with H9N2 or PR8 (MOI=1). (I) Co-IP analysis of the interaction of MAVS with ASB3 and its truncation mutants in HEK-293T cells.

### ASB3 mediates K48-linked polyubiquitination degradation of MAVS

Although prior studies have not identified ASB3 regulation of MAVS protein stability directly, we next sought to determine whether ASB3 affects MAVS expression. We co-transfected HA-tagged ASB3 together with Flag-tagged MAVS, TBK1, or IRF3 and performed immunoblot analysis. We observed that ASB3 inhibited MAVS protein expression but not TBK1 and IRF3 (Fig 5A). To further investigate the mechanisms responsible for ASB3 degradation of MAVS, we treated HEK-293T cells with inhibitors for various protein degradation pathways. We found that ASB3-mediated MAVS degradation could be mostly restored by treatment with the proteasome inhibitor MG132. However, not the autophagy inhibitors 3-methylamine (3-MA), lysosome inhibitor ammonium chloride (NH_4_Cl), or the caspase inhibitor ZVAD (Fig. 5B). Protein ubiquitination is a key step in the ubiquitin-proteasome degradation pathway. Different Immunoblot analyses of K48-linked or K63-linked ubiquitin demonstrated that ASB3 induced polyubiquitination of MAVS by K48-mediated linkage but not K63- (Fig. 5C). Overexpression of ASB3 markedly promoted K48-linked polyubiquitination of MAVS with or without MG132 treatment (Fig. 5D-F). To confirm these results, we used the ASB3 truncates in subsequent experiments. We observed a significant reduction in MAVS polyubiquitination when the SOCX and ANK domains of ASB3 were deleted (Fig. 5G and H). Moreover, overexpression of these truncates of ASB3 suggested that the ANK and SOCX do not suppress the activation of the IFN-β luciferase reporter triggered by MAVS or SeV (Fig. 5I and J). Overall, These data suggest that ASB3 promotes K48-linked polyubiquitination and degradation of MAVS by the proteasome pathway.

**Fig. 5.**
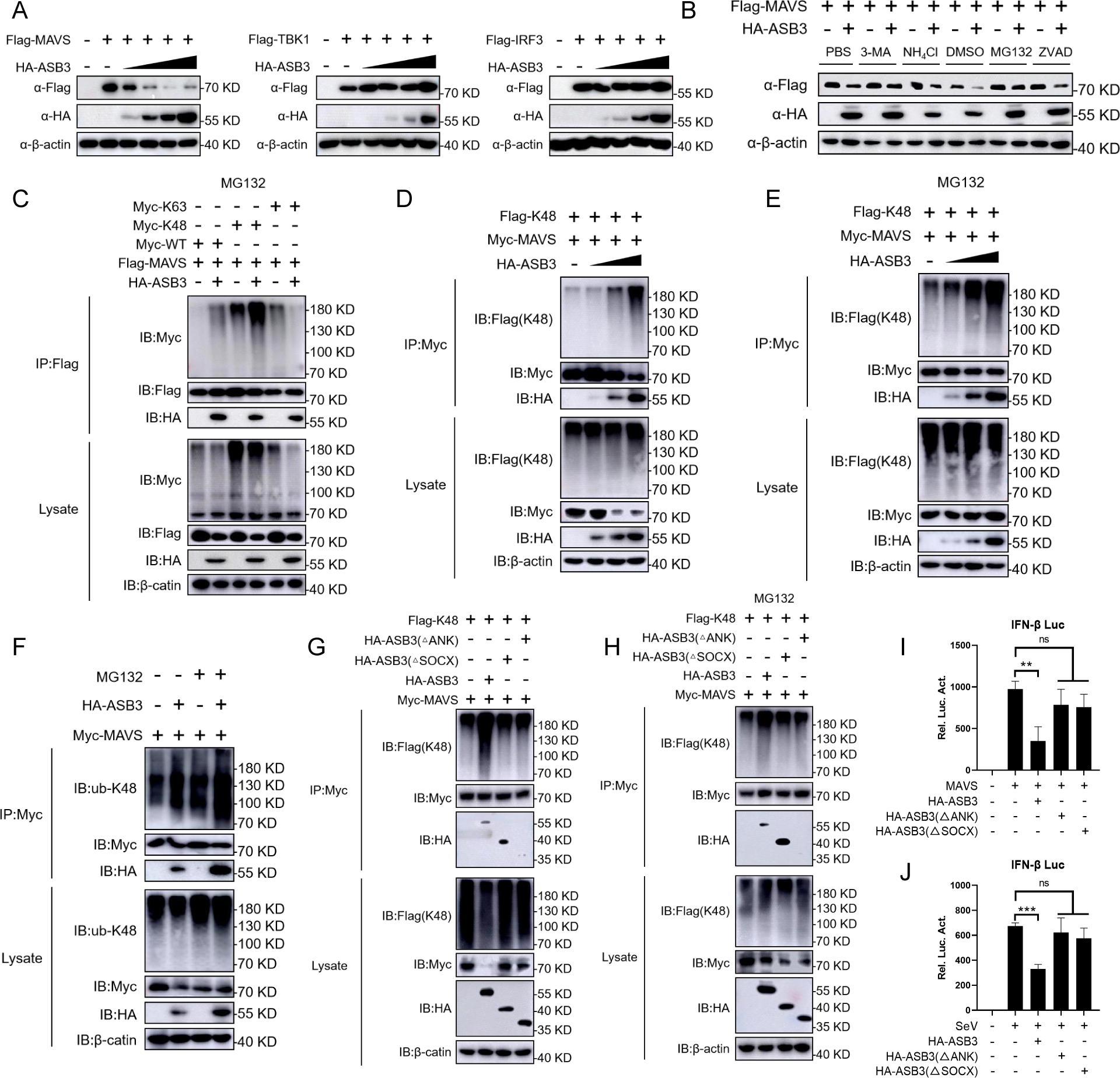
ASB3 potentiates K48-linked polyubiquitination and degradation of MAVS. (A) Immunoblot analysis of HEK-293T cells was transfected with indicated plasmids (MAVS, TBK1, and IRF3) and different concentrations of ASB3 plasmids. (B) HEK-293T cells were transfected with the indicated plasmids for 20 h and then treated with 3-methyladenine (3-MA) (10 mM), NH_4_Cl (20 mM), MG132 (10 μM), or ZVAD (20 μM) for 6 h. The cell lysates were then analyzed by immunoblotting with the indicated antibodies. (C) Co-IP analysis of the polyubiquitination of MAVS in HEK-293T cells transfected with Flag-MAVS, Myc-Ub, or its mutants K48 and K63 [K at indicated residue, and K at other residues were simultaneously mutated to arginines], and ASB3 plasmids and treated with MG132. (D and E) Co-IP analysis of the polyubiquitination of MAVS in HEK-293T cells transfected with Flag-MAVS, Myc-Ub, or its mutants K48, and ASB3 plasmids, and treated with or without MG132. (F) Semi-endogenous Co-IP analysis of the polyubiquitination of MAVS in HEK-293T cells transfected with Flag-MAVS and ASB3 plasmids and treated with or without MG132. (G and H) Co-IP analysis of the polyubiquitination of MAVS in HEK-293T cells transfected with Flag-MAVS, ASB3, or its truncation mutants plasmids, and treated with or without MG132. (I and J) Luciferase reporter assays analyzing IFN-β promoter activity of HEK-293T cells transfected with HA-ASB3 and its deletions along with Flag-MAVS (I) or infected with SeV (J) (MOI=1).

### The Lysine 297 residues of MAVS are critical for ASB3-mediated degradation

To further understand the detailed molecular mechanisms underlying ASB3 function in the degradation of MAVS, we mutated a series of lysines (K) of MAVS and examined their effects on ASB3-mediated MAVS degradation. We found that only mutation of Lysine 297 to Arginine (K297R) abolished ASB3-mediated degradation of MAVS (Fig. 6A). Next, we supplemented various MAVS mutants (K136R, K297R, and K362R) into MAVS^-/-^ HEK-293T cells for further validation. Results from reporter assays showed that of the three loci in close proximity, the MAVS K297R restored ASB3-mediated degradation, which again induced activation of the IFN-β promoter (Fig. 6B). Similarly, MAVS K297R-mediated phosphorylation levels of IRF3 and IκBα could not be downregulated (Fig. 6C). Infection experiments showed that transient overexpression of ASB3 promoted H9N2 or PR8 replication expressing MAVS-WT, MAVS-K136R, and MAVS-K362R (but not K297R) in MAVS^-/-^ HEK-293T cells (Fig. 6D and E). To further analyze the ubiquitination of MAVS at K297 residues, MAVS-WT, MAVS-K136R, or MAVS-K362R was strongly ubiquitinated when myc-tagged K48 ubiquitin was expressed or not, whereas ubiquitination of MAVS-K297R was reduced (Fig. 6F-H). These data indicate that the Lysine 297 residues of MAVS are critical for ASB3-mediated degradation during H9N2 or PR8 infection.

**Fig. 6.**
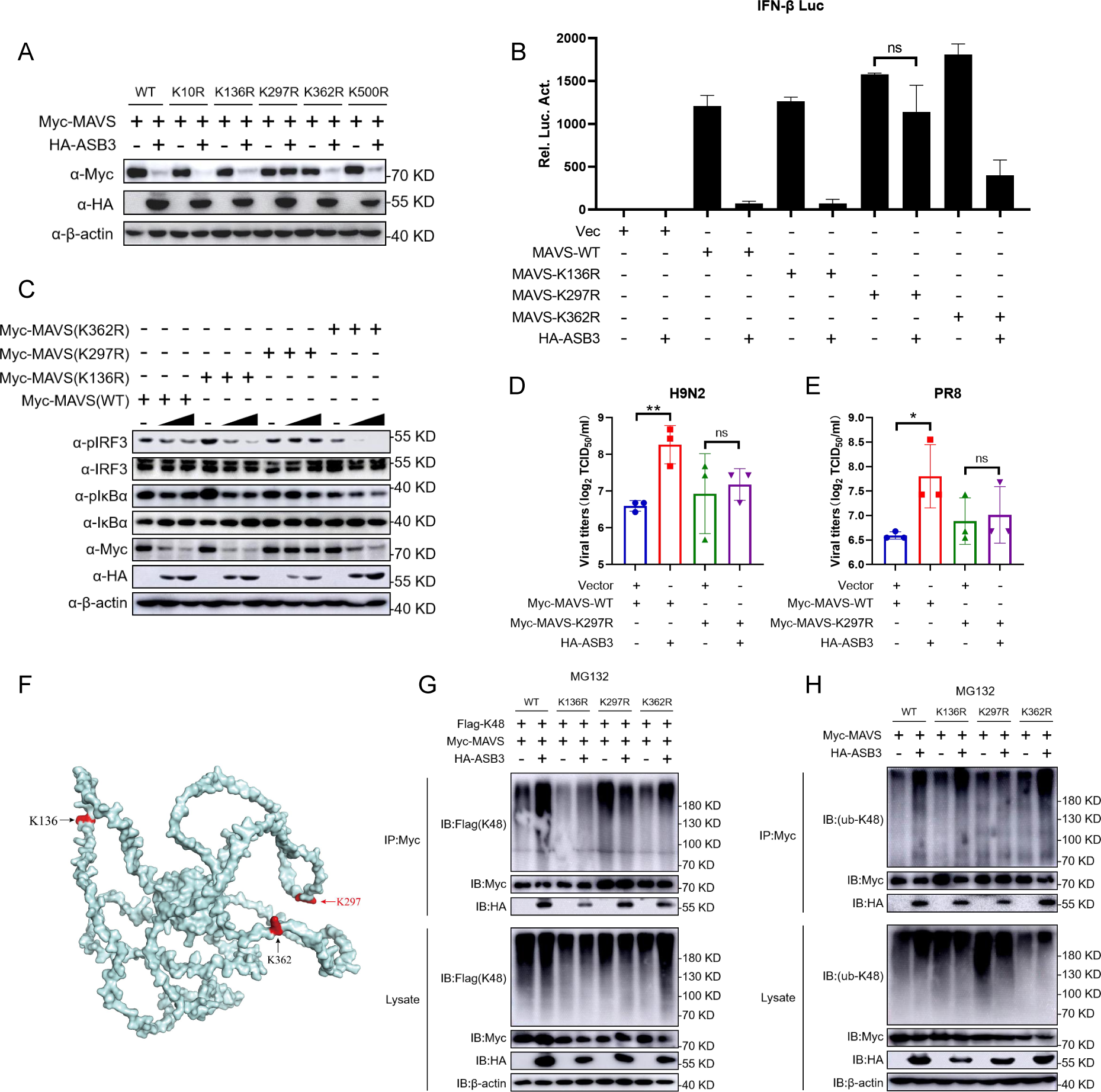
ASB3-mediated MAVS degradation at residues K297. (A) Immunoblots analysis of HEK-293T cells transfected with MAVS or its mutants and ASB3plasmids. (B) The IFN-β and ISRE promoter activation analysis of MAVS^-/-^ HEK-293T cells transfected with MAVS or its mutants and ASB3 plasmids. (C) MAVS^-/-^ HEK-293T cells were transfected with MAVS or its mutants and ASB3 plasmids. The protein expression of pIRF3, IRF3, pIκBα, IκBα, Myc, HA, and β-actin were measured through immunoblot analysis. (D and E) Viral titers in MAVS^-/-^ HEK-293T cells transfected with MAVS or its mutants and ASB3 plasmids and infected with H9N2 virus (D) or PR8 virus (E) (n=3). (F) Molecular model of the MAVS domain generated by PyMOL (PDB:3J6C). (G) Co-IP analysis of the polyubiquitination of MAVS in HEK-293T cells transfected with Myc-MAVS or its truncation mutants, Myc-K48, and HA-ASB3 plasmids, and treated with MG132. (H) Co-IP analysis of the polyubiquitination of MAVS in HEK-293T cells transfected with Flag-MAVS and ASB3 plasmids and treated with or without MG132.

### ASB3 deficiency protects against H9N2 and PR8 infection in mice

To examine the role of ASB3 on influenza virus infection *in vivo*, ASB3^+/+^ mice and ASB3^-/-^ mice were infected with H9N2 and PR8. ASB3^-/-^ mice exhibited a lower rate of body weight loss and a higher survival rate compared to ASB3^+/+^ mice, suggesting that ASB3 deficiency is more resistant to H9N2 and PR8 infection (Fig. 7A and B) (Fig. 7D and E). We also noticed a significant reduction in H9N2 and PR8 titers in the lungs of ASB3^-/-^ mice infected with H9N2 or PR8 on d 5 (Fig. 7C and F). Furthermore, H9N2 or PR8 infection induced more severe histopathological changes in the lungs of ASB3^+/+^ mice. We also observed a more inflammatory cell infiltration in the lungs of ASB3^+/+^ mice, particularly on the 5 d of infection (Fig. 7G-I). Altogether, these results demonstrated that ASB3 deficiency significantly restricts the virulence of H9N2 and PR8 in mice and mitigates the pathological injury caused by viral infection.

**Fig. 7.**
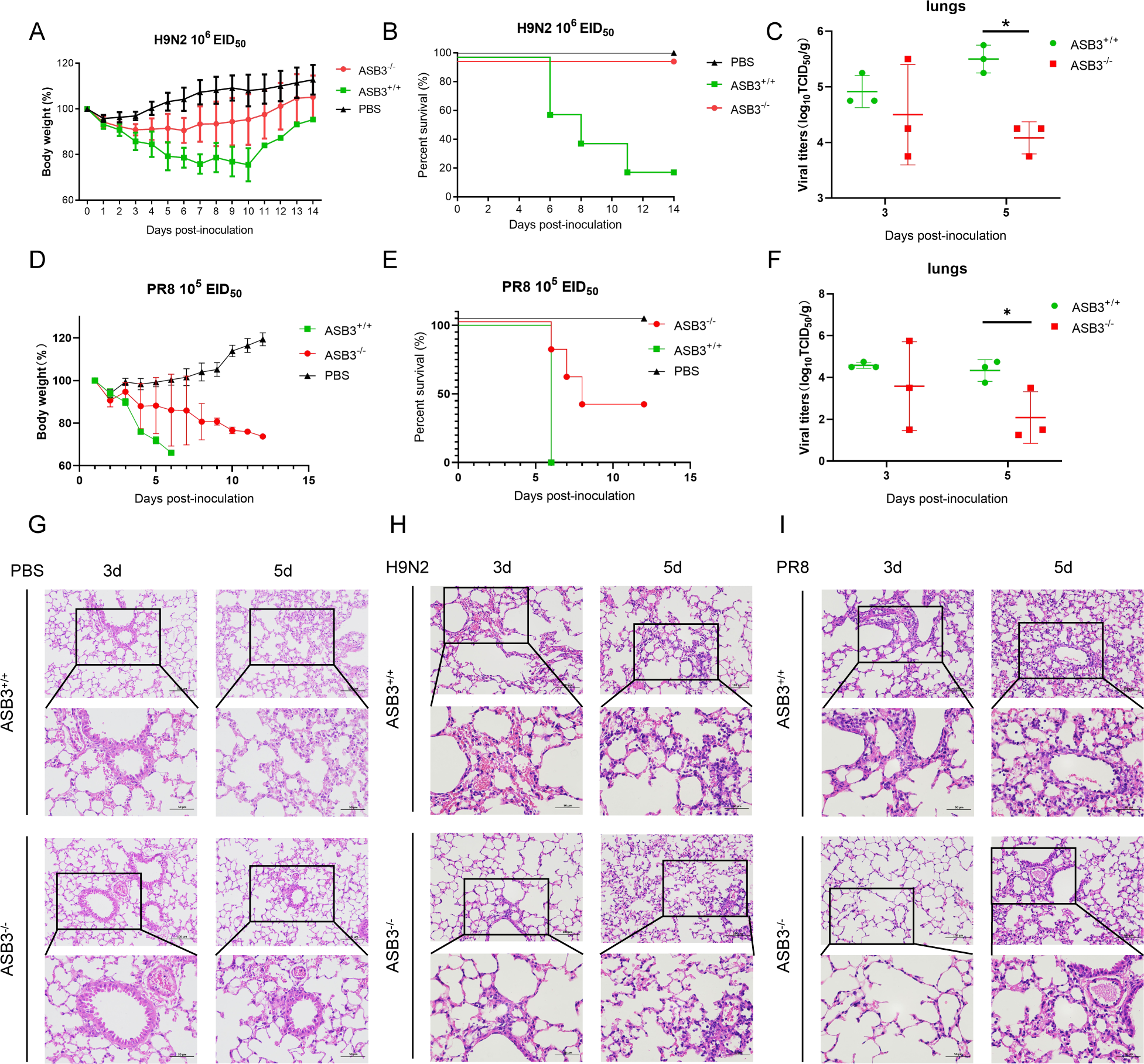
Loss of ASB3 enhances in vivo antiviral immunity. (A and B) Weight loss rate and survival of ASB3^+/+^ and ASB3^-/-^ mice (n = 7 mice/group) infected with H9N2 (1 × 10^6^ PFU/mouse) via intranasal infection. (C) TCID_50_ assay was used to test H9N2 titer in mice’s lungs. (D and E) Weight loss rate and survival of ASB3^+/+^ and ASB3^-/-^ mice (n = 7 mice/group) infected with PR8 (1 × 10^5^ PFU/mouse) via intranasal infection. (F) TCID_50_ assay was used to test PR8 titer in mice’s lungs. (G-I) Representative H&E-stained images of lung sections from ASB3^+/+^ and ASB3^-/-^ mice infected with H9N2 and PR8 for 3 d and 5 d. Scale bars, 50 or 100 µm.

### ASB3 negatively regulates *in vivo* antiviral responses by degrading MAVS

To further clarify the role of ASB3 in inhibiting IFN-I production *in vivo*, we performed phosphorylation analyses on lung tissues from ASB3^+/+^ and ASB3^-/-^ mice infected with H9N2 and PR8. We found that ASB3 deficiency enhanced the phosphorylation levels of TBK1, IRF3, and IκBα in the lungs. At the same time, ASB3 exacerbated the degradation of MAVS during infection. In contrast, MAVS expression levels in the lungs of ASB3^-/-^ mice returned to normal expression levels on d 5. Notably, lower expression levels of NP were detected in ASB3-deficient lung tissues (Fig. 8A and B). qPCR analysis demonstrated that ASB3 deficiency also enhanced the gene transcription levels of IFN-β1 and ISG15 in the lungs (Fig. 8C and D). These data all indicate that ASB3 deletion effectively upregulates the IFN-I response during influenza virus infection. To further corroborate the association between ASB3 and MAVS during *in vivo* infection, we conducted co-IP and ubiquitination analyses. The results showed that ASB3 strongly interacts with MAVS after infection with H9N2 and PR8 (Fig. 8E) . Consistent with these results, ASB3 enhanced K48-linked polyubiquitination of MAVS in the lungs of mice infected for 5 d, whereas ASB3 deficiency weakened such phenomenon (Fig. 8F). Taken together, our findings suggest that ASB3-promoted degradation of MAVS inhibits the antiviral immune response induced by H9N2 and PR8 infection.

**Fig. 8.**
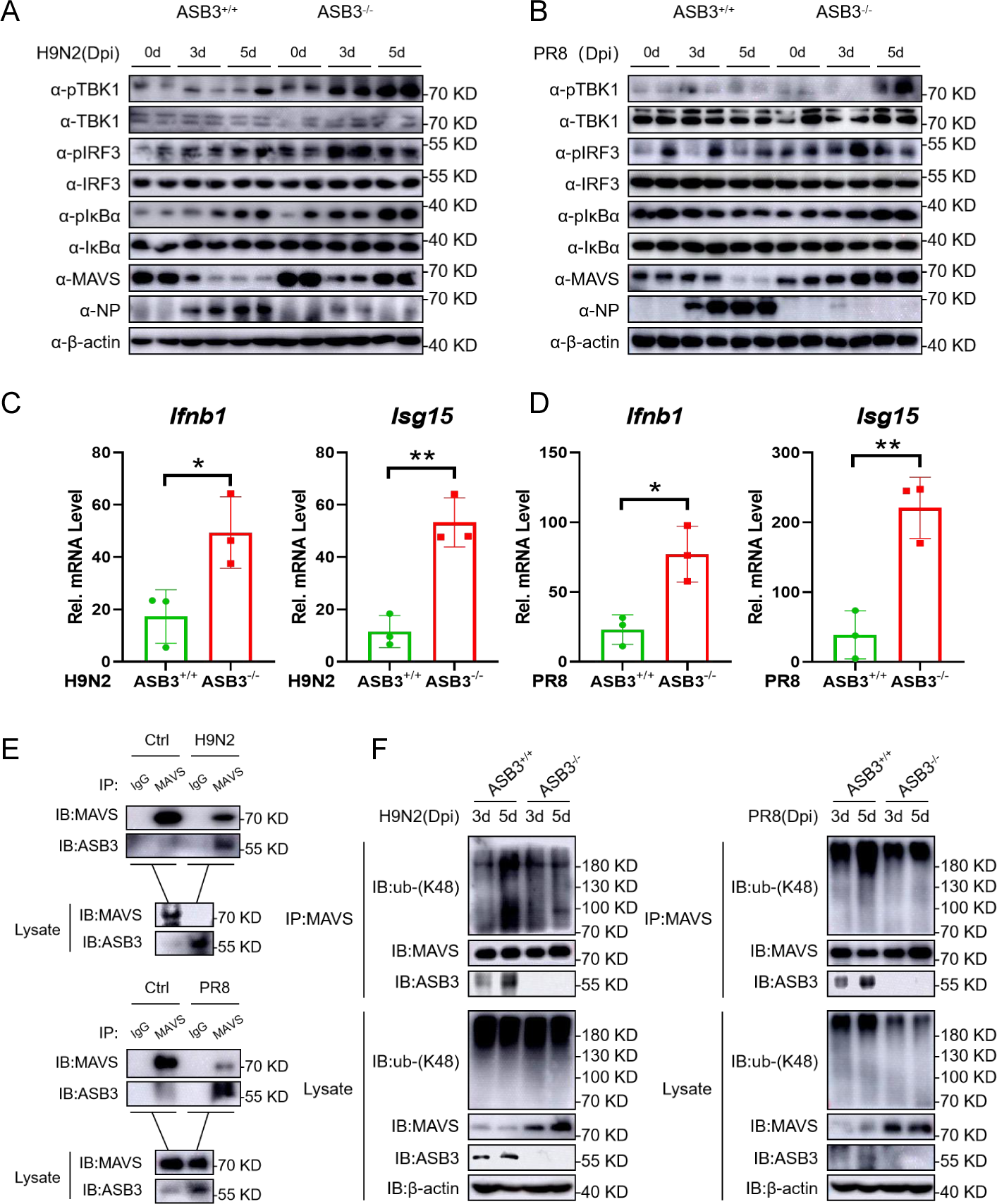
ASB3 deficiency eliminates MAVS degradation in the lungs. (A and B) Immunoblot analysis of pTBK1, TBK1, pIRF3, IRF3, pIκBα, IκBα, MAVS, NP, and β-actin expression in the lungs from the ASB3^+/+^ and ASB3^-/-^ mice infected with H9N2 or PR8 for 3 d and 5 d. (C and D) qPCR analysis of Ifnb1 and Isg15 mRNA expression in the lungs from the ASB3^+/+^ and ASB3^-/-^ mice infected with H9N2 or PR8 for 3 d and 5 d. (E) Co-IP analysis of the interaction of MAVS with ASB3 in the lungs from the ASB3^+/+^ mice infected with H9N2 or PR8 for 5 d. (F) Co-IP analysis of the polyubiquitination of MAVS in the lungs from the ASB3^+/+^ and ASB3^-/-^ mice infected with H9N2 or PR8 for 3 d and 5 d.

## DISCUSSION

Following an extended period of co-evolution and competition, hosts have evolved a number of potent defense mechanisms against viral infections. However, an increasing number of host suppressors that target the negative regulatory functions of the antiviral immune system are being identified. In the present study, we demonstrate that ASB3 inhibits antiviral innate immunity by showing that overexpression of ASB3 impairs SeV- and H9N2-induced activation of IFN-I, and ASB3-deficient mice are more resistant to H9N2 and PR8 infection. Moreover, during H9N2 and PR8 infection, ASB3 loss increases MAVS expression levels and IRF3 phosphorylation. Interestingly, ASB3 catalyzes K48-linked polyubiquitination of lysine residues in MAVS, which reduces IFN-I signaling (Fig. 9).

**Fig. 9.**
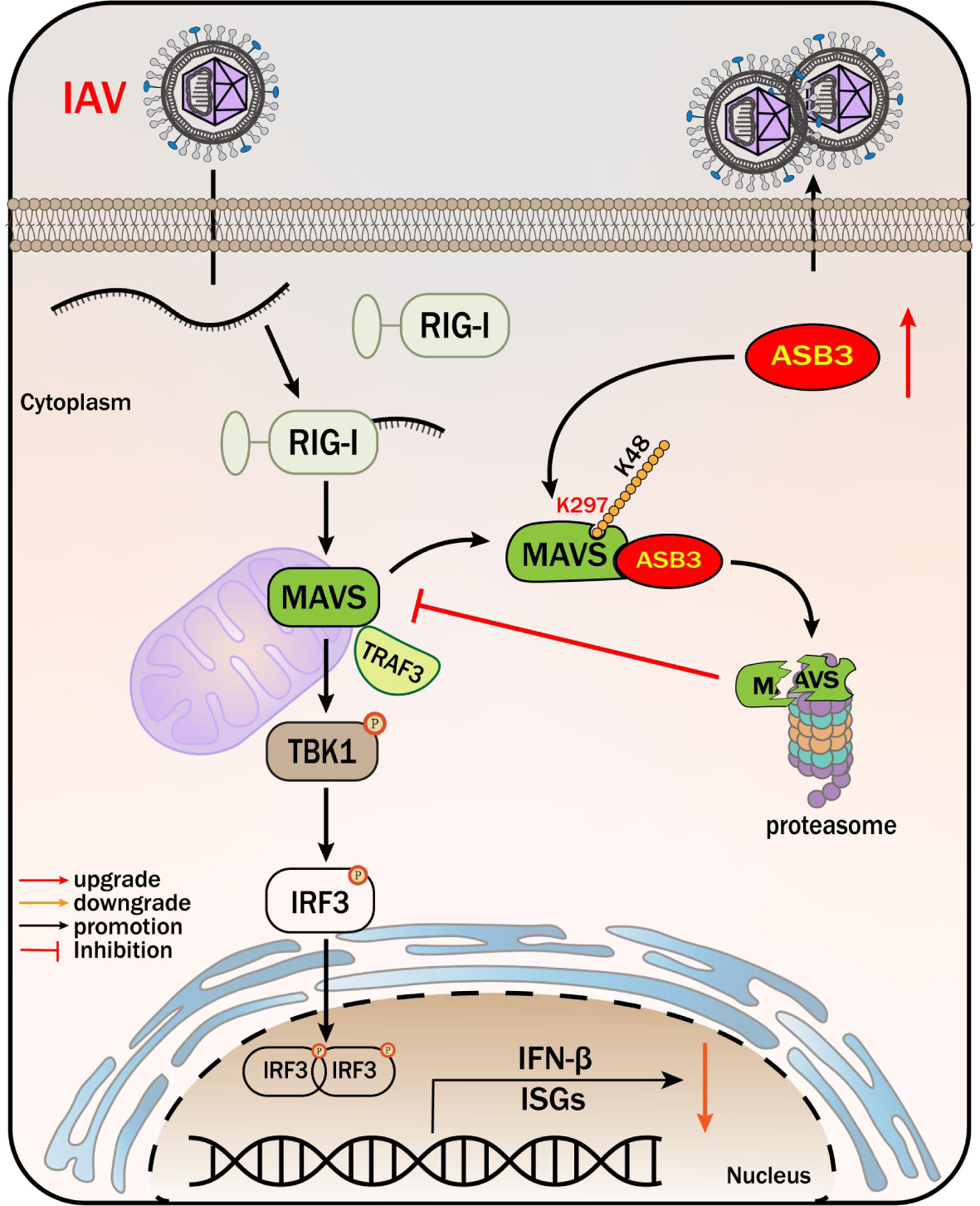
A working model of ASB3 inhibition of antiviral immunity. Upon RNA virus infection, ASB3 expression is induced to be upregulated. ASB3 associates with MAVS and mediates its K48-linked polyubiquitination at the K297. This modification blocked the association of MAVS and TBK1, which inhibited the phosphorylation of TBK1 and IRF3, thereby impairing the transcriptional activation of IFN-I.

Viral nucleic acids are sensed by RIG-I and cGAS, which then trigger the IFN-β signaling cascade and hundreds of ISGs to be produced. Our study revealed that ASB3 expression may be enhanced in response to RNA virus infection (Fig. 1A). Overexpression of ASB3 inhibited SeV- and poly (I:C)-induced IFN-β production (Fig. 2A), whereas ASB3 deletion mediated elevation of IFN-I (Fig. 3). Numerous malignancies have been linked to the ASB family of proteins, and a variety of tumor types show abnormal expression of ASB3 (20). According to one study, ASB3 prevents CRC metastases by lowering the expression of the Vimentin protein (21). Notably, recent research has revealed that Vimentin proteins act as negative regulators of antiviral immunity during RNA and DNA viral infections (22). In contrast to these findings, our study uncovers a novel biological function of ASB3.

The C-terminal structural domain of MAVS allows it to bind to the outer mitochondrial membrane, where it localizes. As a crucial adapter protein for RLR signaling, it plays a pivotal role in mediating signaling pathways (23). Upon viral infection, MAVS needs to maintain a delicate balance between activation and inactivation to ensure a host antiviral immune response of appropriate intensity. In our study, we confirmed that ASB3 interacts specifically with MAVS (Fig. 4C-H).

Crucially, we also discovered that ASB3 is inactive in relation to TRAF3, cGAS, and STING (Fig. 4D), which explains the unique response of ASB3 in RNA virus infection. Similarly, another study found that by interacting with MAVS, the E3 ubiquitin ligase RNF115 efficiently counteracts excessive antiviral responses (24).

The overreaction of host proteins is mostly dependent on the strategy of escape immunization of viral proteins. Previous studies have shown that the host protein phosphatase PPM1G dephosphorylates MAVS, thereby inhibiting MAVS activation (25).

Furthermore, we showed that MAVS is degraded by ASB3 in a dose-dependent manner (Fig. 5A), and this occurs via the ubiquitin-proteasome pathway. (Fig. 5B). As the major route for removing damaged proteins, the ubiquitin-proteasome system (UPS) is essential for preserving intracellular homeostasis. Several emerging classes of host inhibitory factors that target the functions of MAVS have been identified. For instance, GP73 promotes MAVS degradation through a proteasome-dependent pathway (26). Recent studies have claimed that E3 ligase Smurf1 enhances proteasome-mediated MAVS degradation (27). Actually, ubiquitination is a major process influencing the post-translational modification (PTM) of proteins. We confirmed that ASB3-triggered K48-linked polyubiquitination of MAVS inhibited the activation of IFN-I signaling (Fig. 6C-F). Subsequently, by mapping, we found that ASB3 mediated the K48-linked polyubiquitination of MAVS at Lys297. (Fig. 7E-G). Lys48 (K48)-linked and Lys63 (K63)-linked polyubiquitination are well recognized as the two most common types of polyubiquitination in ubiquitin-mediated protein modification. Certain E3 enzymes are responsible for conferring substrate specificity. Previous research has shown that the E3 ligase MARCH3 can facilitate K48-linked polyubiquitination at IL-3Rα K377, which has the effect of suppressing inflammatory responses that are generated by IL-3 (28). Furthermore, it has been reported that the E3 ligase TRIP catalyzes K48-linked polyubiquitination repair of TBK1 at Lys372, promoting autophagy-lysosomal degradation of TBK1(29). According to a recent study, RIG-I-mediated MAVS aggregation is initiated by Riplet, a novel RIG-I-independent MAVS E3 ligase, through the catalysis of unanchored K63-linked polyubiquitin chains (30). Naturally, there have been reports of other types of ubiquitination modifications possessing immunomodulatory functions. Huang et al. found that PKR is polyubiquitinated in a K33-linked manner by the E3 ligase HECTD3, with PKR’s Lys68 serving as the main ubiquitination site for the HECTD3-modified innate response to RNA virus infection (31).

Importantly, ASB3^-/-^ mice reduced susceptibility to influenza A virus (Fig. 7). We have observed before that ASB3 deletion does not activate or suppress innate immune signaling in the absence of viral infection. However, under certain pathological circumstances, ASB3 is overactivated to exert its immunosuppressive function (32). We also observed reduced H9N2 and PR8 titers in the lungs of ASB3^-/-^ mice (Fig. 7C and F), which we attribute to an IFN-I antiviral immune response limited by ASB3-mediated impairment of MAVS function (Fig. 8). E3 enzymes can also target viral proteins to achieve limitation or aid in viral reproduction, as multiple investigations have demonstrated. Yuan et al. found that E3 enzyme RNF5-mediated a K63-linked polyubiquitination modification at the Lys15 of the M protein promoted SARS-CoV-2 replication (33), and Liu et al.reported that MARCH8 inhibited IAV release by redirecting the viral M2 protein from the plasma membrane to the lysosome for degradation (34). More research is required to determine whether or not ASB3 aids in the production or release of viral proteins.

To sum up, we first proposed that the E3 ligase ASB3, whose expression is upregulated in influenza A virus infection, negatively regulates the innate immune response mediated by RLRs. Our study identifies a mechanism by which ASB3 controls the antiviral defense. The knowledge gained from these investigations will help develop ASB3 as a possible target for medications that combat influenza viruses.

## MATERIALS AND METHODS

### Mice

ASB3^-/-^ mice were generated using PiggyBac transposon-based targeting technology and were kindly provided by Prof. Dongqin Yang (Shanghai Fudan University). Mice used in this study were housed under specific-pathogen-free (SPF) conditions and had an FVB genetic background. ASB3^+/+^, ASB3^+/-^, and ASB3^-/-^ mice were identified by PCR amplification and DNA sequencing with the primers

5′-CTGAGATGTCCTAAATGCACAGCG-3′,

5′-CCATGACCAAACCCAATTTACACAC-3′, and

5′-TTGTTTTTTTTCCCCCCTAGACAGG-3′, respectively.

### Cells, viruses, and plasmids

Human embryonic kidney cells (HEK-293T), Madin-Darbey Canine Kidne cells (MDCK) and human lung epithelial cells (A549) were cultured in Dulbecco’s modified Eagle’s medium (DMEM) or DMEM/F12 supplemented with 10% (v/v) fetal bovine serum (FBS) and 2 mM glutamine with penicillin (100 U/ml)/streptomycin (100 mg/ml). BMDM and primary kidney epithelial cells were dissociated from the bone marrow and kidney of WT or ASB3^-/-^ mice. Isolated BMDM were resuspended in RPMI-1640 medium, counted, and cultivated in growth medium with 10% (v/v) FBS, 2 mM glutamine, penicillin (100 U/ml)/streptomycin (100 mg/ml), 10 mM HEPES, 1 mM sodium pyruvate, and 20 ng/ml recombinant M-CSF for 5 d (35). MAVS^-/-^ cells were stored in our laboratory. All cells were cultured and maintained at 37°C with 5% CO_2_.

A/chicken/Hong Kong/G9/1997 (H9N2) and A/Puerto Rico/8/34 (H1N1, PR8) were kindly provided by Prof. Wentao Yang (Jilin Agricultural University, China). Sendai virus (SeV), Recombinant vesicular stomatitis virus expressing green fluorescent protein (VSV-GFP), and herpes simplex virus 1 (HSV-1) were stored in our laboratory (36).

Plasmids for Flag-tagged RIG-IN, MAVS, TRAF3, cGAS, STING, TBK1, IRF3, IRF3-5D, ubiquitin (Flag-K48), HA-tagged MDA5-N, Myc-tagged ubiquitin (Myc-WT, Myc-K48, and Myc-K63), MAVS mutants (WT, K136R, K297R, and K362R), IFN-β-Luc, ISRE-Luc, and pRL-TK (internal control luciferase reporter plasmid) used in the study were described previously (10). Plasmids for HA-tagged ASB3, ASB3 (ΔANK), and ASB3 (ΔSOCX) were kindly provided by Prof. Dongqin Yang (Fudan University, Shanghai, China).

### Viral infections

Cells were grown to approximately 70% confluence and infected with H9N2, PR8, SeV, VSV-GFP, and HSV-1. Cells were exposed to either virus dilutions supplemented with or without TPCK trypsin, incubated at 37°C and 5% CO_2_for 2 h, and supernatants were then removed. Cell monolayers were rinsed with PBS to remove unattached virus particles and then incubated in the virus maintenance fluid (2% FBS) at 37°C and 5% CO_2_ for the designated time.

### Antibodies and reagents

The antibodies used in this study were as follows: HRP-conjugated anti-HA (12013819001), Myc (11814150001) antibodies (Roche); phosphorylated IκBα (AF5851), IκBα (AG2737), HRP-labeled Goat anti-Mouse IgG(H+L) (A0216), and HRP-labeled Goat anti-Rabbit IgG(H+L) (A0208) antibodies (Beyotime), FITC Goat Anti-Mouse IgG (H+L) (K1201), Cy5 Goat Anti-Rabbit IgG (H+L) (K1212) secondary antibodies (APExBIO); anti-HA (66006-2-Ig), anti-DYKDDDDK (20543-1-AP), anti-β-actin (66009-1-Ig), and anti-TRAF3 (18099-1-AP) antibodies (Proteintech); anti-mus MAVS antibody (E8Z7M), anti-ubiquitin (K48) (8081) and HRP-conjugated mouse anti-rabbit IgG (Conformation Specific) (5127) antibodies (Cell Signaling Technology); anti-ubiquitin (WT) antibody (Santa Cruz, sc-8017); anti-ASB3 (88812) antibody (MBL); HRP-conjugated anti-Flag (A8592) antibody (Sigma); anti-Hu MAVS/VISA antibody (A300-782A) (BETHYL). Reagents used in the study included: 3-Methyladenine (M9281), MG132 (M7449), DMSO (D2650), NH_4_Cl (09718), anti-Myc agarose affinity beads (A7470) and protein A/G agarose affinity beads (P6486/E3403) (Sigma); mouse recombinant M-CSF (315–02) (PeproTech); DAPI (C1005), ZVAD (Caspase inhibitor Z-VAD-FMK, C1202), NP-40 (ST366), HEPES Solution (C0215) (Beyotime). TnT® T7/SP6 Quick Coupled Transcription/Translation System (L1170/L2080) (Promega).

### Coimmunoprecipitation (Co-IP) and Immunoblotting analysis

HEK-293T cells at 80-90% confluency were washed with PBS three times and collected into tubes using a cell scraper. The same mass of lung tissue was washed three times with precooled PBS and mechanically homogenized with RIPA lysis buffer (Thermo Fisher) containing 1% PMSF. All samples were lysed in NP-40 lysis buffer (20 mM Tris-HCl, 1 mM EDTA, 1% NP-40, and 150 mM NaCl) supplemented with Halt Protease Inhibitor Cocktail and kept on ice for 30 min. After 30 min of incubation, the lysates were spun down at 12000 rpm at 4°C for 20 min. All supernatants were collected, and protein was quantified using the BCA Protein Assay Kit (Beyotime). The supernatants were pretreated with 30 µl of anti-Flag agarose affinity gels or protein A/G at 4°C for 2 h. The indicated primary antibody or IgG control (mouse IgG, Beyotime) was then added to the pretreated lysates and incubated overnight at 4°C. The IP complexes and whole-cell lysates were subjected to protein transfer by the conventional Western blotting method, and protein development was detected using the antibodies mentioned above. Immunoblots were observed using Amey Imager 600RGB and quantified using ImageJ.

### Dual-luciferase reporter assays

Plasmids encoding Flag-tagged RIG-IN, MAVS, cGAS, STING, TBK1, and IRF3-5D or HA-tagged ASB3 were co-transfected with IFN-β-Luc, ISRE-Luc, and pRL-TK into HEK-293T cells for 24 h using Lipofectamine 3000 (Invitrogen) reagent. Cell samples were collected at the indicated times and lysed or assayed using the Dual-Luciferase® Reporter Gene (DLR™) Assay System (Promega) according to the manufacturer’s instructions. Finally, the relevant reporter gene activity is assayed with Firefly luciferase and Renilla luciferase reagents.

### Quantitative real-time RT-PCR (qPCR) analysis

Total RNA was isolated from the transfected or infected cells using TRIzol (Thermo Fisher). 2 μg RNA was used to synthesize first-strand cDNA using Moloney mouse leukemia virus (M-MLV) (Promega). The amplification of specific PCR products was detected using SYBR Premix Ex TaqTM II (Vazyme). Real-time PCR was conducted using the Applied Biosystems 7500 real-time PCR System. The results were displayed as relative expression values normalized to GAPDH.

### Histology

Lungs from H9N2 or PR8-infected or uninfected mice were dissected, fixed in 4% paraformaldehyde solution overnight, embedded into paraffin, sectioned, stained with hematoxylin and eosin solution, then examined by microscope (Leica) for histological changes.

### Confocal microscopy

HEK-293T or A549 cells were collected at the indicated times. Cells were fixed with 4% paraformaldehyde for 15 min, then permeabilized and blocked in containing 0.5% Triton X-100 and 5% skimmed milk for 1 h. Cells were stained with specific primary antibodies and then blotted with fluorescence-coupled secondary antibodies. Cell nuclei were labeled with DAPI. The stained cells were observed and photographed with a Zeiss microscope (LSM 710).

### Virus titration

Viral titers of virus-containing culture supernatants were determined by end-point titration. For tissue infection samples, each sample was serially diluted 10-fold, and for cell infection samples, each sample was serially diluted 2-fold. The diluted samples were inoculated into MDCK cells. Supernatants were collected 48 hours after inoculation and assayed for chicken erythrocyte agglutination ability as an indicator of a virus. Replicated infectious virus titers were reported as log10 TCID_50_/ml or log2 TCID_50_/ml and were calculated from three replicates using Reed-Muench’s method.

### Statistical analysis

Data are presented as the mean ± SD unless otherwise indicated. All samples were analyzed using GraphPad Prism 8 software. Data were analyzed by unpaired two-tailed Student’s *t*-test or two-way ANOVA with Sidak’s test to correct for multiple comparisons. Probability (p) values of < 0.05 were considered significant: \**P* < 0.05, \*\**P* < 0.01, \*\*\**P* < 0.001, and \*\*\*\**P* < 0.0001; n.s., not significant.

