## Supplementary material for "E3 ligase ASB3 downregulates antiviral innate immunity by targeting MAVS for ubiquitin-proteasomal degradation": Table. 1

**Table 1 The primer sequences for q-PCR.**

| **Primers** | **Sequence (5′→3′)** |
| --- | --- |
| *Hu GAPDH*-Forward | AAAATCAAGTGGGGCGATGCT |
| *Hu GAPDH*-Reverse | GGGCAGAGATGATGACCCTTT |
| *Hu IFNB1*-Forward | CAGCAATTTTCAGTGTCAGCAAGCT |
| *Hu IFNB1*-Reverse | TCATCCTGTCCTTGAGGCAGTAT |
| *Hu IFIT2*-Forward | AAGCACCTCAAAGGGCAAAAC |
| *Hu IFIT2*-Reverse | TCGGCCCATGTGATAGTAGAC |
| *Hu ISG15*-Forward | AGGACAGGGTCCCCCTTGCC |
| *Hu ISG15*-Reverse | CCTCCAGCCCGCTCACTTGC |
| *Hu OASL*-Forward | CTGATGCAGGAACTGTATAGCAC |
| *Hu OASL*-Reverse | CACAGCGTCTAGCACCTCTT |
| *Mus GAPDH*-Forward | ACGGCCGCATCTTCTTGTGCA |
| *Mus GAPDH*-Reverse | ACGGCCAAATCCGTTCACACC |
| *Mus Ifnb1*-Forward | TCCTGCTGTGCTTCTCCACCACA |
| *Mus Ifnb1*-Reverse | AAGTCCGCCCTGTAGGTGAGGTT |
| *Mus Isg15*-Forward | GGTGTCCGTGACTAACTCCAT |
| *Mus Isg15*-Reverse | CTGTACCACTAGCATCACTGTG |
| *Mus Cxcl10*-Forward | CCAAGTGCTGCCGTCATTTTC |
| *Mus Cxcl10*-Reverse | GGCTCGCAGGGATGATTTCAA |
| *Mus Ccl5*-Forward | GCTGCTTTGCCTACCTCTCC |
| *Mus Ccl5*-Reverse | TCGAGTGACAAACACGACTGC |
| *Mus Mx-1*-Forward | GACCATAGGGGTCTTGACCAA |
| *Mus Mx-1*-Reverse | AGACTTGCTCTTTCTGAAAAGCC |
| *Mus Oasl1*-Forward | CAGGAGCTGTACGGCTTCC |
| *Mus Oasl1*-Reverse | CCTACCTTGAGTACCTTGAGCAC |
| *Mus Isg56*-Forward | CTGAGATGTCACTTCACATGGAA |
| *Mus Isg56*-Reverse | GTGCATCCCCAATGGGTTCT |
| *Mus IL-6*-Forward | TCTGCAAGAGACTTCCATCCAGTTGC |
| *Mus IL-6*-Reverse | AGCCTCCGACTTGTGAAGTGGT |
| *Mus TNFα*-Forward | TCTTCTCATTCCTGCTTGTGG |
| *Mus TNFα*-Reverse | GGTCTGGGCCATAGAACTGA |
